# Molecular mechanisms underlying enhanced hemichannel function of a cataract-associated Cx50 mutant

**DOI:** 10.1101/2021.08.18.456706

**Authors:** J.J. Tong, U. Khan, B.G. Haddad, P.J. Minogue, E.C. Beyer, V.M. Berthoud, S.L. Reichow, L. Ebihara

**Author notes:** Indicates equal contribution (J.J. Tong, U. Khan).

## Abstract

Connexin-50 (Cx50) is among the most frequently mutated genes associated with congenital cataracts. While most of these disease-linked variants cause loss-of-function due to misfolding or aberrant trafficking, others directly alter channel properties. The mechanistic bases for such functional defects are mostly unknown. We investigated the functional and structural properties of a cataract-linked mutant, Cx50T39R (T39R), in the *Xenopus* oocyte system. T39R exhibited greatly enhanced hemichannel currents with altered voltage-gating properties compared to Cx50 and induced cell death. Co-expression of mutant T39R with wild-type Cx50 (to mimic the heterozygous state) resulted in hemichannel currents whose properties were indistinguishable from those induced by T39R alone, suggesting that the mutant had a dominant effect. Co-expression with Cx46 also produced channels with altered voltage-gating properties, particularly at negative potentials. All-atom molecular dynamics simulations indicate that the R39 substitution can form multiple electrostatic salt-bridge interactions between neighboring subunits that could stabilize the open-state conformation of the N-terminal domain, while also neutralizing the voltage-sensing residue D3 as well as residue E42 which participates in loop-gating. Together, these results suggest T39R acts as a dominant gain-of-function mutation that produces leaky hemichannels that may cause cytotoxicity in the lens and lead to development of cataracts.

**Statement of significance:** We investigated the functional and structural properties of a cataract-linked mutant, Cx50T39R (T39R), in the *Xenopus* oocyte system and showed that T39R exhibited greatly enhanced hemichannel currents with altered voltage-gating properties compared to Cx50 and induced cell death. Consistent with our experimental findings, all-atom equilibrium state molecular dynamics (MD) simulations of T39R show that R39 stabilized the open-state configuration of the N-terminal (NT) domain from an adjacent subunit. These results suggest that T39R causes disease by preventing the hemichannels from closing when present in the plasma membrane in the undocked state and provide an atomistic rationalization for the Cx50 disease-linked phenotype. They also expand our understanding of how connexin hemichannel channel gating is controlled.

## Introduction

The eye lens is an avascular, transparent organ whose main function is to focus light on the retina to support vision. It is composed of epithelial cells and fiber cells that are interconnected by a vast network of gap junctions. These gap junctions allow the flow of fluid, ions and nutrients throughout the lens with minimal blocking of light transmission compared to a circulation system dependent on blood vessels, which would block and distort the transmission of light (see [1] for a review). Gap junctions are composed of many intercellular channels, each of which is composed of two connexons or hemichannels (residing in the plasma membrane of neighboring cells) that dock to form a complete gap junctional channel. Hemichannels can also exist as undocked channels which form large, relatively nonselective conductances in single plasma membranes that are gated by external calcium and transmembrane voltage [2, 3].

Gap junctional channels in vertebrates are formed by a family of closely related proteins called connexins, with 21 members in humans [4]. Three different types of gap junctional proteins that show a differential pattern of expression have been identified in the lens: Cx43 [5], Cx50 [6] and Cx46 [7]. Both Cx50 and Cx46 are highly expressed in lens fiber cells, where they may form heteromeric channels [8, 9], while Cx43 is expressed primarily in epithelial cells together with Cx50. The importance of these proteins in the lens has been confirmed by studies which showed that genetic ablation of either Cx50 or Cx46 led to the development of cataract in mice [10, 11]. Moreover, mutations in Cx50 and Cx46 have been linked to congenital cataract in humans and mice (see [12] for a review). Indeed, there are more cataract-associated Cx50 variants than for any other gene expressed in the lens [13].

When expressed in heterologous expression systems, most of these cataract-linked connexin mutations are associated with a decrease or loss of gap junctional communication, either because of biosynthetic defects or degradation that decrease the abundance of connexins at the plasma membrane, or because the mutants form non-functional gap junctional channels [14–19]. Some of the mutants also act as dominant negative inhibitors of co-expressed wild-type lens connexin function [15, 16, 20]. At least one of the mutations causes a gain-of-hemichannel function [17, 21].

In the present study, we examined the functional properties of the Cx50 mutation, T39R (replacement of threonine by arginine at amino acid position 39), that has been associated with congenital cataract [22, 23]. We show that this mutant causes a dominant gain-of-hemichannel function when expressed in *Xenopus* oocytes. Consistent with our experimental findings, all-atom equilibrium state molecular dynamics (MD) simulations of T39R, based on the high-resolution cryo-EM structure of Cx50 in the open-state [24], shows that R39 stabilizes the open-state configuration of the N-terminal (NT) domain from an adjacent subunit. These results suggest that mutation of T39R causes disease by preventing the hemichannels from closing when present in the plasma membrane in the undocked state. This would cause cell depolarization and the release of intracellular metabolites into the extracellular spaces, ultimately resulting in cell death. These results further support and extend our understanding of the functional roles played by key residues within Cx50 that contribute to channel gating properties.

## Materials and Methods

### Generation of constructs

Mutations were introduced into human Cx50 cDNA in a SP64T vector with QuikChange Site-Directed Mutagenesis Kit (Stratagene) by using PCR primers that amplified the whole plasmid. The primers used were as follows: T39R, sense 5′-ctc atc ctt ggc agg gcc gca gag ttc-3′ and antisense 5′-gaa ctc tgc ggc cct gcc aag gat gag-3′; T39A, sense 5′-ctc atc ctt ggc gcg gcc gca gag-3′ and antisense 5′-ctc tgc ggc cgc gcc aag gat gag-3′; and T39S, sense 5′-cct cat cct tgg ctc ggc cgc aga g-3′ and antisense 5′-ctc tgc ggc cga gcc aag gat gag g-3′. The coding regions of all amplified constructs were fully sequenced at the Cancer Research Center DNA Sequencing Facility of the University of Chicago (Chicago, IL) to confirm the absence of additional unwanted mutations.

### Expression of connexins in Xenopus oocytes

Connexin cRNAs were synthesized using the mMessage mMachine *in vitro* transcription kit (Ambion, Austin, TX) according to the manufacturer’s instructions. The amount of cRNA was quantitated by measuring the absorbance at 260 nm.

Adult female *Xenopus* laevis frogs were anesthetized with tricaine and a partial ovariectomy was performed in accordance with protocols approved by the Rosalind Franklin University Animal Care and Use Committee. The oocytes were manually defolliculated after treating them with collagenase type 4 (Worthington Biochemical, Lakewood, NJ). Stage V and VI oocytes were selected. For some of the experiments, single, collagenase-treated oocytes (Ecocyte Biosciences, Austin, TX) were used. The oocytes were pressure-injected using a Nanoject variable microinjection apparatus (model mo. 3-000-203, Drummond Scientific, Broomal, PA) with 0.05 – 500 ng/μl of connexin cRNA along with 5 ng/36.8 nl of oligonucleotide antisense to mRNA for *Xenopus* Cx38 to prevent contamination by endogenous connexin hemichannel currents [25]. The total volume of injected cRNA and oligonucleotides was held constant at 36.8 nl in all of the experiments. The oocytes were incubated for 16 to 72 hours at 18°C in Modified Barth’s Solution (MBS) containing 0.7 mM CaCl_2_ prior to performing the electrophysiological experiments. Only those oocytes that had very low levels of endogenous connexin hemichannel currents were used for data collection and analysis.

### Immunoblotting

Oocytes were injected with 2 ng/µl T39R, T39A, T39S or wild type Cx50 cRNA + Cx38 antisense oligonucleotides, or antisense alone and incubated for 24 hours in MBS containing 0.7 mM [Ca^2+^]_o_ at 18°C. Healthy oocytes were fastfrozen in liquid nitrogen and stored at -80°C. Oocytes were homogenized in 1 ml of homogenization buffer (5 mM Tris · HCl, 1 mM EDTA, 2 mM EGTA, 2 mM phenylmethylsulfonyl fluoride, pH 8.0, containing cOmplete EDTA-free protease inhibitor cocktail at a concentration of 1 tablet/7 ml (Roche Applied Science, Indianapolis, IN, USA)) by repeated passage through a 20-gauge needle to obtain plasma membrane-enriched preparations. Homogenates were then centrifuged at 3,000 g for 5 min at 4°C to pellet yolk granules. The supernatant was then centrifuged at 100,000 g for 1 hour at 4°C on a Sorvall RC M120EX microultracentrifuge, and the pellet was resuspended in the homogenization buffer as previously described [6, 26]. Proteins were resolved by SDS-PAGE on 8% acrylamide gels and electrotransferred onto Immobilon P (Millipore, Bedford, MA). Then, the membranes were stained with Ponceau S (to verify equal electrotransfer of the proteins) before being subjected to immunoblotting as previously described [21, 27]. The X-ray films were scanned on a flat-bed scanner (Epson Perfection V700 Photo; Epson America, Long Beach, CA) to obtain a digital image on which to quantify the bands. Their density was quantified using Adobe Photoshop CS3 (Adobe Systems Inc., San Jose, CA) as previously described [28]. The results are reported in arbitrary units. Graphs were prepared using SigmaPlot 10.0 (Systat Software, Inc.).

### Oocyte viability experiments

For the oocyte viability experiments, the oocytes were injected with 2 ng/µl of T39R or Cx50 cRNA and incubated in MBS either in the nominal absence of external calcium or in the presence of 3 mM [Ca^2+^]_o_ for 48 hours at 18°C. The oocytes were scored for cell death using cell blebbing and disorganization of pigment in the animal pole as markers at 19, 22, 24, 43 and 48 hours after injection.

### Electrophysiology

Hemichannel currents were recorded from single oocytes using a GeneClamp 200 two-electrode voltage-clamp amplifier (Molecular Devices, Sunnyvale, CA) as previously described [29]. The standard external bath solution contained (in mM) 88 choline Cl or 88 NaCl, 1 KCl, 2.4 NaHCO_3_, 1 MgCl_2_, and 15 HEPES, pH 7.4 to which different concentrations of calcium were added.

The normalized conductance-voltage curves were determined from tail currents as previously described [29]. Briefly, the membrane potential was stepped from −10 mV to the test potential for 20 s and then hyperpolarized to −80 mV. Initial tail current amplitudes at -80 mV were determined by fitting the tail currents to a single exponential or the sum of 2 exponentials and then extrapolating back to t = 0. The initial tail currents were normalized to the initial tail current amplitude after a 40 mV test pulse and plotted as a function of test potential.

Pulse generation and data acquisition were performed using a PC computer equipped with PCLAMP9 software and a Digidata 1322A data acquisition system (Molecular Devices). The currents were sampled at 2 kHz and low pass filtered at 50-100 Hz. Leak correction, when applied, was performed by subtracting the average leakage current in control oocytes (oocytes injected only with AS). All experiments were performed at room temperature (20 – 22° C).

Data analysis and graphing were performed using PCLAMP10.6 software (Molecular Devices), OriginPro 2021 (OriginLab, Northampton, Massachusetts), and SigmaPlot 14 (Systat software, San Jose, California). Group statistics were reported as mean ± SEM.

### Molecular dynamics simulation and analysis

Visual Molecular Dynamics (VMD) v.1.9.3 [30] was used to build Cx50 (PDB: 7JJP) [24] mutant systems for molecular dynamics (MD) simulations. Side chains were protonated according to neutral conditions, and the protonated HSD model was used for all histidine residues. Disulfide bonds identified in the experimental structures were enforced for all models. Amino acids corresponding to the cytoplasmic loop (CL) connecting TM2 and TM3 (residues 110 – 148) and the C-terminal domain (residues 237 – 440) were not included, as experimental data describing the structure of these large flexible regions are unresolved. The N- and C-terminal residues resulting from the missing CL (R109 and R149) were neutralized. N-terminal acetylation was added at the G2 position (G2_ACE_) in VMD through an all-atom acetylation patch using the AutoPSF plugin, to mimic the *in-vivo* co-translational modification identified in the native protein[9, 31].

The T39R mutation was introduced using two separate techniques, to allow comparison of different starting conformations. VMD’s *mutator* plugin [30] was used to create a naïve model, where arginine was placed in the exact same conformation as the threonine it replaced (Model 1). SWISS-MODE [32] was used on the Cx50 template (PDB: 7JJP) to generate a mutant homology model, with a minimized initial conformation (Model 2).

Both resulting structures were then embedded in 1-palmitoyl-2-oleoyl-sn-glycero-3-phosphocholine (POPC) lipid bilayers generated by CHARMM-GUI [33] using VMD’s *mergestructs* plugin. Lipids that overlapped with the protein and the pore were removed, and the systems were placed in water boxes using VMD’s *solvate* plugin. Water that overlapped with the lipid bilayers was then removed. Hexagonal boundary conditions were used, with circumradius 70 Å and height 200 Å. The systems were then neutralized using the *autoionize* plugin, followed by the addition of 150 mM KCl and 150 mM NaCl to the solvent areas corresponding to intracellular and extracellular regions of the simulation box, respectively. Finally, hydrogen mass repartitioning [34] was applied to both systems to enable a 4 fs timestep.

Nanoscale Molecular Dynamics (NAMD) v.3.0 alpha [35] was used for all classical molecular dynamics (MD) simulations, using the CHARMM36 force field [36] for all atoms and TIP3P explicit solvent model. Periodic boundary conditions were used to allow for the particle mesh Ewald calculation of electrostatics [37]. Both systems were prepared following the same minimization and equilibration protocol, as follows. First, the lipid tails were allowed to minimize with all other atoms fixed for 1 ns using a 1 fs timestep, allowing the acyl chains to “melt” with constant volume at a temperature of 300 K (NVT). All subsequent simulations were performed using the Langevin piston Nosé-Hoover method for pressure control (NPT). Next, the entire system, including lipids, solvent, and ions, was allowed to minimize with the protein harmonically constrained (1 kcal mol^-1^), for 2 ns using a 2 fs timestep. Another 2 ns minimization step was then applied, also with a 2 fs timestep, in which the system was free to minimize with a harmonic constraint applied only to the protein backbone (1 kcal mol^-1^), to ensure stable quaternary structure while side chains relax in their local environment. The entire system was then released from all constraints and subject to all-atom equilibration using a Langevin thermostat (damping coefficient of 0.5 ps^-1^), with a constant temperature of 310 K and constant pressure of 1 atm, using a 4 fs timestep for 30 ns. Finally, both systems were simulated in triplicate for 100 ns production runs, again with a 4 fs timestep. Each replica (n = 3) started from the end of a 30 ns equilibration phase with velocities reinitialized.

Root mean squared deviations (r.m.s.d.), comparing the backbone conformations of MD simulation to the original starting structures, and root mean square fluctuations (r.m.s.f.), comparing the amplitudes of backbone fluctuations during MD simulation, were calculated using VMD. Both T39R models approached a steady r.m.s.d. during the equilibration phase and maintained stability during all production runs, and r.m.s.f. values were comparable to previous simulations of Cx50 [24].

To assess interactions introduced by the R39 mutation, we recorded which residues were within 4 Å of R39 every 10 ps, and aggregated counts across chains and replicates. Once key residues were identified, the distances between each R39 and the residues of interest were recorded every 10 ps across all replicates of all production runs. The points of reference for R39 (C_ζ_), D3 (C_ɣ_), and E42 (C_δ_) were chosen to capture equivalent rotameric states. Since the acetylated G2 contains two carbonyl oxygen atoms, either of which can interact with R39, the minimum of the distances between R39 and the oxygen atoms was recorded as the distance to G2_ACE_. Histograms (bin size ≈ 0.1 Å) and corresponding empirical distribution functions (ECDFs) were then plotted.

Based on the distance histograms, 4.5 Å was chosen as the cutoff distance below which R39 and another residue are considered to be interacting. This criterion was used to generate time series of binary interaction states for each R39 and each contact residue, recorded every 10ps. Dwell times were obtained by calculating the length of consecutive interaction states, ignoring periods of noninteraction that lasted less than 500 ps. Any interactions with a final length of less than 1 ns were ignored. Histograms (bin size = 4 ns) and corresponding ECDFs were then plotted.

Nonbonding energies between residue 39 and G2_ACE_ or D3 were computed across all trajectories using the *namdenergy* plugin, with a 1 ns timestep. Each chain was treated independently from all replicates (n = 36 for T39R models and n = 24 for Cx50) to calculate the final summary statistics, plotted as mean ± SEM. A two-sample independent t-test was used to compare sample means to determine statistical significance.

## Results

### Functional characterization of Cx50 T39R hemichannels

In our initial experiments, we observed that expression of T39R in *Xenopus* oocytes appeared to have a toxic effect on oocytes. To study this phenomenon in more detail, we injected oocytes with 2 ng/µl of T39R or Cx50 cRNA and incubated them in Modified Barth’s Solution in the presence or absence of external calcium. In the absence of external calcium, most of the T39R cRNA-injected oocytes developed cell blebbing, cytoplasm leakage, and pigmentation changes consistent with cell death by 24 hours after injection, as illustrated in Figure 1A. In contrast, Cx50 cRNA injected oocytes and control oocytes injected only with antisense oligonucleotides to the endogenous *Xenopus* Cx38 (AS) remained healthy without any obvious changes in membrane pigmentation. The survival time of the T39R expressing oocytes was significantly prolonged by incubating the oocytes in external medium containing 3 mM [Ca^2+^]_o_. The results of 4 independent experiments are summarized in Figure 1B. Less than 5% of the control or Cx50-injected oocytes died by 48 hours, regardless of the absence or presence of added calcium. In contrast, 98% of the T39R cRNA-injected oocytes died by 48 hours when incubated in the absence of calcium and 38% died in the presence of calcium.

**Figure 1.**
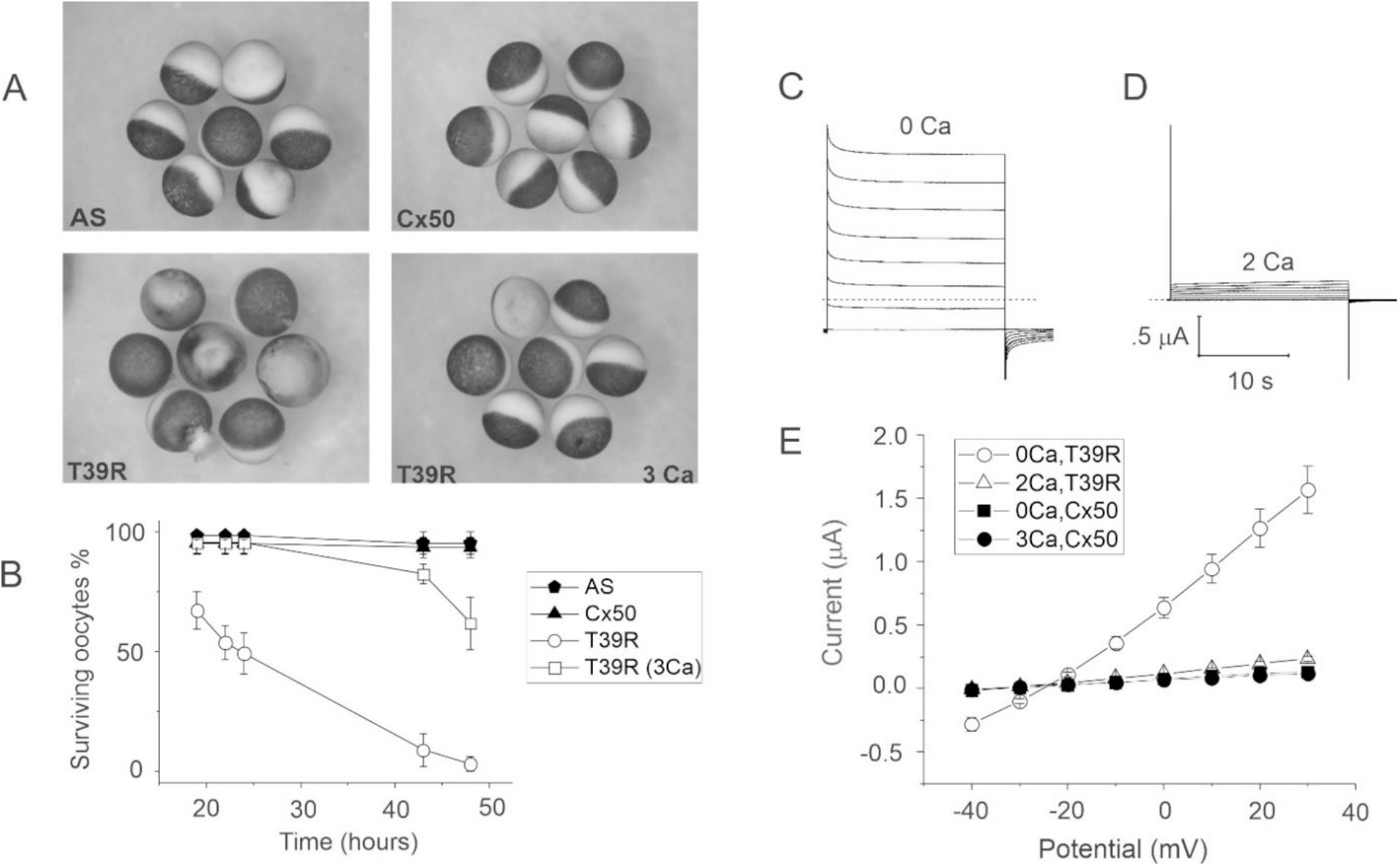
T39R has a toxic effect on oocytes that can be rescued by increasing external calcium. A. Oocytes injected with T39R cRNA show obvious signs of cell death (cell blebbing and pigment disorganization in the animal pole) after a 24 hour incubation period in modified Barth’s solution (MBS) containing zero-added calcium (bottom left). In contrast, Cx50 cRNA-injected oocytes (Cx50) and antisense-injected control oocytes (AS) remained healthy when incubated under identical conditions (top left and right). Raising the external calcium from 0 to 3 mM rescued the T39R cRNA-injected oocytes from cell death (bottom right). B. Graph shows the percentage of surviving oocytes/total number of oocytes at various times following injection of 2 ng/µl wild-type or mutant cRNA. The total number of oocytes in each group ranged between 14-17. Each point in panel B represents the mean ± SEM of 4 independent experiments. C and D. Families of current traces recorded from a T39R cRNA-injected oocyte before and after application of 2 mM calcium, respectively. The voltage clamp protocol consisted of a series of sequential steps from a holding potential of -40 mV to 30 mV in 10 mV increments. In the presence of nominally zero calcium, a large hemichannel current was observed that was mostly activated at a holding potential of -40 mV. Application of 2 mM [Ca^2+^]_o_ caused a marked reduction in the amplitude of the current at both positive and negative potentials. The dashed line represents zero current level. E. Steady-state I-V relationships (measured at the end of the pulse) in Cx50 (closed symbols) and T39R (open symbols) cRNA-injected oocytes before and after application of calcium. Each point in panel E represents the mean ± SEM of 5-6 oocytes. All of the data were obtained with oocytes from a single donor frog. Similar results were obtained with oocytes from at least 3 other donors.

One explanation for these findings is enhanced hemichannel activity of T39R channels that is partially blocked by increasing external calcium. To investigate this possibility, we injected oocytes with extremely low concentrations of mutant cRNA (0.05 – 0.25 ng/µl) that were insufficient to induce cell death and we studied the oocytes at times ranging between 18 – 72 hours after injection using a two-electrode-voltage-clamp technique. Oocytes injected with 0.25 ng/µl of T39R cRNA exhibited large hemichannel currents that were partially blocked by raising [Ca^2+^]_o_ from nominally zero to 2 mM and had a reversal potential of ∼-25 mV in choline chloride external solution, as illustrated in Figure 1C-E. In contrast, oocytes injected with 3 ng/µl Cx50 cRNA or antisense-injected control oocytes showed little or no calcium-sensitive hemichannel currents, consistent with the results of previous studies [17, 21].

### Functional comparison of T39R, T39S and T39A variants

To further examine the role of residue 39 in Cx50 hemichannel gating, threonine 39 was replaced by alanine (the amino acid at the equivalent position in the first transmembrane spanning domain of Cx46) or serine (another hydroxyl group-containing amino acid). T39A and T39S cRNA-injected oocytes developed hemichannel currents whose size was much greater than that of Cx50, but significantly lower than that observed in T39R cRNA-injected oocytes. Therefore, to facilitate comparison of voltage gating properties of the three mutants, we injected oocytes with 10-30 times lower amounts of T39R cRNA to reduce the size of its hemichannel currents to levels similar to those observed for T39A and T39S.

To compare levels of expression of the different Cx50 constructs, we performed immunoblot analysis of membrane-enriched samples prepared from oocytes injected with antisense Cx38 oligonucleotides alone or in combination with similar amounts of cRNA encoding T39R, T39S, T39A, or Cx50. Homogenates of oocytes injected with wild-type or mutant cRNAs contained a predominant band with an Mr of ∼61 kDa (Figure 2A). The densitometric value of the bands of the mutant proteins were similar or lower than the densitometric value of the Cx50 band (Figure 2B). No immunoreactive bands were detected in antisense-injected control oocytes. Thus, these results excluded the possibility that increased hemichannel activity could result from increased mutant protein production.

**Figure 2.**
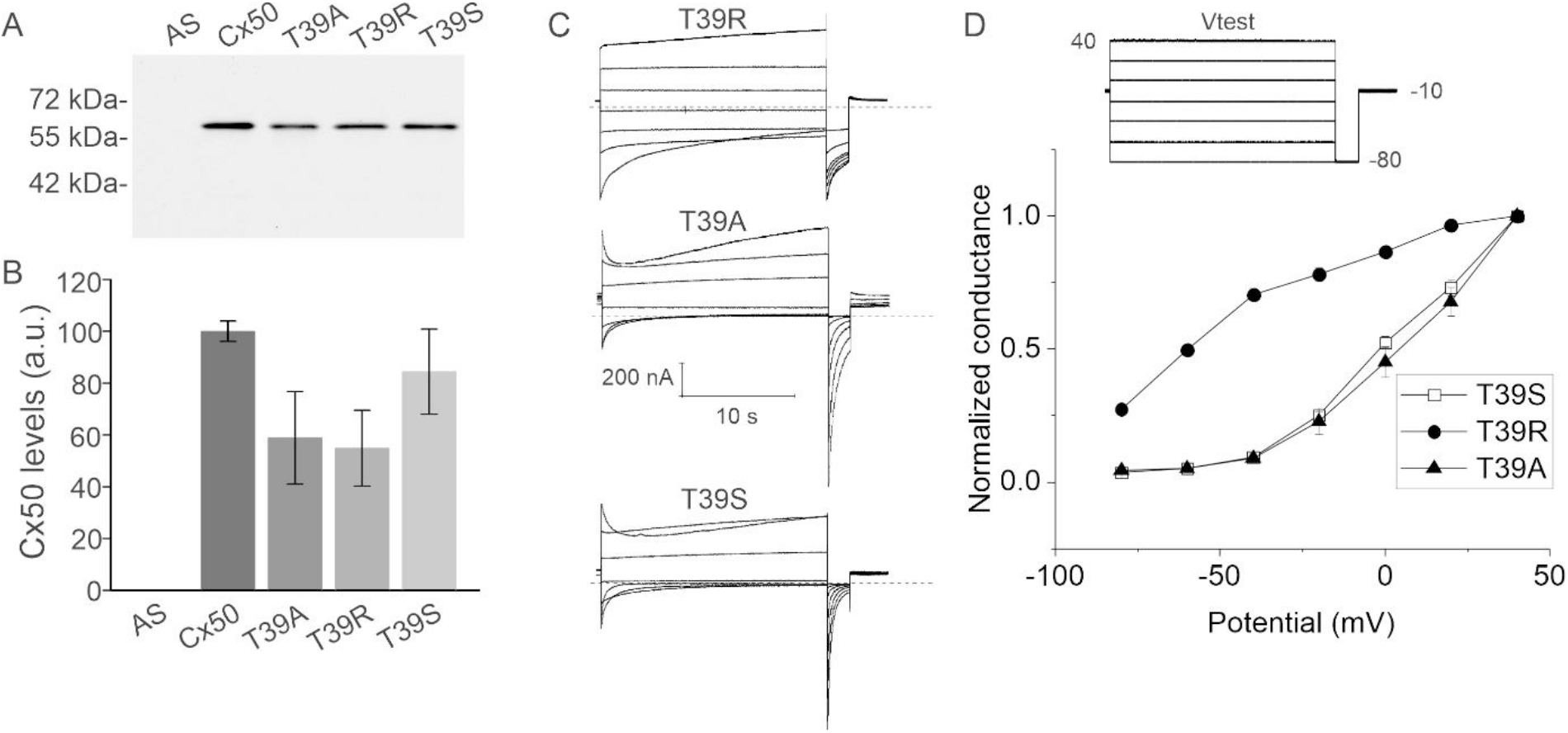
Hemichannel currents formed from T39R, T39S and T39A show differences in V_j_ and loop gating. A. Immunoblot analysis of plasma membrane enriched samples prepared from *Xenopus* oocytes injected with similar amounts of Cx50, T39R, T39S or T39A cRNA in combination with antisense Cx38 oligonucleoties or with antisense Cx38 oligonucleotides alone and incubated for 24 hours at 18° C. B. The graph shows the mean ± SEM of the densitometric values of the immunoreactive bands obtained from 4 or 5 biological replicates expressed in arbitrary units (a.u.). The mean densitometric value of the band for Cx50 was similar or higher than the mean densitometric values of the bands for the mutants indicating that the protein levels of Cx50 were similar or higher than those of oocytes expressing T39R, T39S or T39A. C. Representative families of membrane current traces recorded from single oocytes injected with cRNA for T39R (top), T39A (middle) or T39S (bottom) in sodium chloride solution containing 0.7 mM [Ca^2+^]_o_ using the pulse protocol shown in Figure 2D (inset). The dashed lines represent zero current level. The data were corrected for leakage. D. Normalized G-V curves for T39R (solid circles; n = 5), T39A (solid triangles; n = 4), T39S (open squares; n = 6) determined by plotting the amplitude of the tail current at -80 mV normalized to the tail current following a step to +40 mV as a function of test potential. Each point represents the mean ± SEM.

Figure 2C compares the electrophysiological properties of hemichannel currents recorded from single oocytes injected with cRNA for T39R, T39A or T39S in sodium external solutions containing 0.7 mM [Ca^2+^]_o_. The pulse protocol consisted of a 20-s test pulse, ranging between −80 mV and +40 mV from a holding potential of −10 mV, followed by a hyperpolarizing pulse to −80 mV. When the oocytes were bathed in 0.7 mM [Ca^2+^]_o_, all 3 mutant hemichannels exhibited a negative voltage-gating process, termed loop gating [38], that tended to cause the channels to close at negative potentials and slowly open at positive potentials. However, the rate and extent of hemichannel closure at negative potentials was greatly reduced for T39R compared to T39A and T39S. In addition, the normalized G-V curve for T39R, determined by plotting the relative amplitude of the tail current at −80 mV as a function of test potential, was shifted to the left with respect to that observed for T39A and T39S (Figure 2D). At positive potentials, T39A and T39S hemichannels exhibited another voltage-gating process termed V_j_ gating [38], which has been previously shown to close Cx50 hemichannels at positive potentials [39, 40]. This gating process was reduced in T39R hemichannels. Thus, our findings suggest that replacement of T39 by arginine, but not by serine or alanine, results in a loss of voltage-dependent gating at both positive and negative potentials.

### Functional characterization of T39R mixed with Cx50 and Cx46

To mimic the effect of heterozygous expression, we investigated the effect of mixing Cx50 and T39R by injecting oocytes with a 1:1 mixture of Cx50 and T39R. Oocytes co-expressing Cx50 and T39R showed hemichannel currents whose amplitude and voltage dependent gating properties resembled those observed in oocytes injected with T39R cRNA alone, indicating that T39R modified channel gating in a dominant manner (Figure 3A,B).

**Figure 3.**
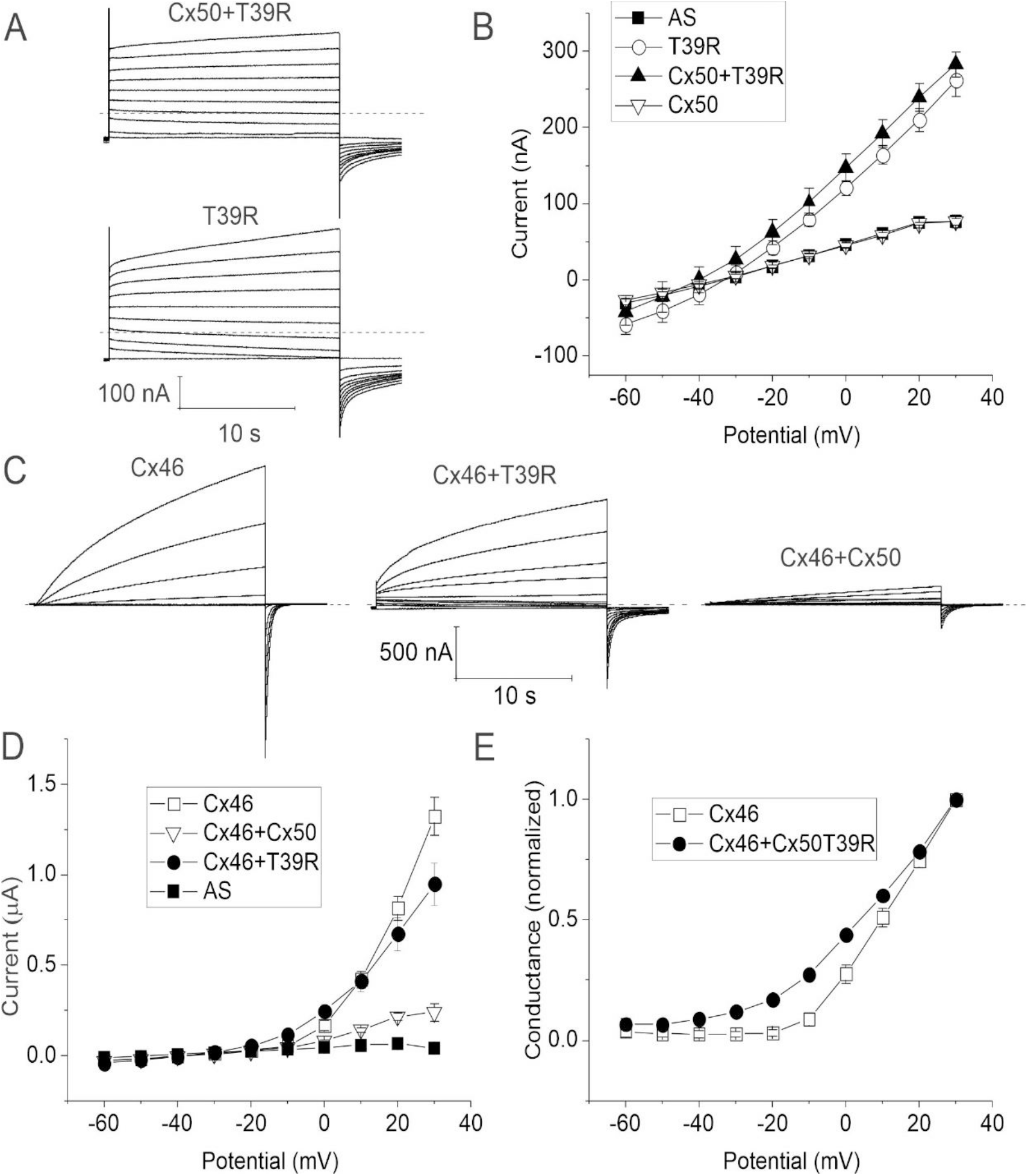
T39R acts as a dominant gain-of-hemichannel function mutation when co-expressed with wild-type Cx50 or Cx46. A. Representative families of current traces recorded from oocytes injected with a 1:1 mixture of T39R+Cx50 cRNA (top) or T39R cRNA alone (bottom) in response to a series of voltage clamp steps applied from a holding potential of -60 mV to potentials between -60 and +30 mV in increments of 10 mV. The amount of T39R cRNA injected into each oocyte was held constant. The oocytes were bathed in NaCl solution containing 0.7 mM [Ca^2+^]_o_. Dashed lines represent zero current. B. Steady-state I-V curves for T39R (open circles, n = 5), T39R+Cx50 (solid triangles, n = 3), Cx50 (open inverted triangles, n = 5) and antisense-treated controls (AS; solid squares, n = 7). Each point represents the mean ± SEM. All data were obtained with oocytes from a single donor frog. Similar results were obtained in the presence of nominally zero [Ca^2+^]_o_ (data not shown). C. Representative families of current traces recorded from oocytes injected with Cx46 cRNA alone (left), a 1:1 mixture of Cx46+T39R cRNA (middle), or a 1:1 mixture of Cx46+Cx50 cRNA (right) in response to voltage clamp steps applied from -60 mV to +30 mV in 10 mV increments from a holding potential of - 60 mV. The amount of Cx46 cRNA injected into each oocyte was held constant. The oocytes were bathed in NaCl solution containing 0.7 mM [Ca^2+^]_o_. The data were corrected for leakage. D. Steady-state I-V curves for Cx46 (open squares, n = 6), Cx46+Cx50 (open inverted triangles, n = 6), Cx46+T39R (solid circles, n = 7), and AS (solid squares, n=4). E. Normalized G-V curves for Cx46 (open squares, n = 5) and Cx46+T39R (solid circles, n = 7). Each data point in panels D and E represents the mean ± SEM. All data were obtained with oocytes from a single donor frog.

Because a second connexin (Cx46) is co-expressed with Cx50 in the lens and shown to be capable of co-assembling into heteromeric channels [8, 9], we studied the effect of mixing of Cx50 or T39R with Cx46 on hemichannel function as shown in Figure 3C-E. As expected, Cx46 formed time- and voltage-dependent currents that slowly activated at membrane potentials more positive than -10 mV. On repolarization to -60 mV, a large, inward tail current was observed that decayed to baseline over a period of several seconds. The main effect of co-expressing Cx50 with Cx46 was to reduce the size of the resulting hemichannel currents compared to those seen in oocytes injected with only Cx46, in agreement with our previously described results [17]. In contrast, oocytes co-injected with a 1:1 mixture of T39R and Cx46 cRNA showed large hemichannel currents, with magnitudes similar to that observed in oocytes injected with Cx46 alone. However, the mixed currents showed a slowing of the time course of deactivation at negative potentials and a negative shift in the threshold for activation compared to Cx46. These findings suggest that both wild-type and mutant Cx50 can interact with Cx46 to form heteromeric channels with altered gating properties.

### MD-simulation of Cx50 T39R

To gain mechanistic insight into the gain-of-function observed by T39R, we conducted all-atom equilibrium-state MD simulation studies, using the previously determined 1.9 Å structure of native sheep Cx50 as a template [24]. Sheep Cx50 contains ∼95% sequence identity (∼98% similarity) to human over the structured domains. T39R models were constructed *in silico* by two independent methods (Model #1 and #2), to allow comparison of different starting conformations (see Methods). In both models, the N-terminal residue G2 is acetylated (G2_ACE_) to match the prominent state found in the lens (Myers et al., 2018). In Cx50, T39 is located in the middle of transmembrane helix 1 (TM1), and positioned toward the hydrophobic anchoring residue W4 on the NT domain (Figure 4A,B). The NT domain is thought to function both as the voltage-sensing domain and as a gate to close the channel [40–43]. Homology modeling of arginine at this site introduces a steric clash with the indole ring of W4 (Figure 4B), which by naïve interpretation might be expected to destabilize the open-state conformation of Cx50. However, following energy minimization and all-atom MD equilibration, R39 is able to reorient and found to adopt a fluctuating range of conformational states that are compatible with the previously characterized open state conformation of the NT domain (Figure 4B) [9]. Indeed, root-mean-square-fluctuation (r.m.s.f.) analyses of the NT domain, which reports on the conformational stability, were indistinguishable between the T39R models and Cx50 in multiple replicate MD simulations (Figure 4C and [24]). These results indicate that T39R does not significantly disrupt the overall architecture or stability of the open-state of the channel.

**Figure 4.**
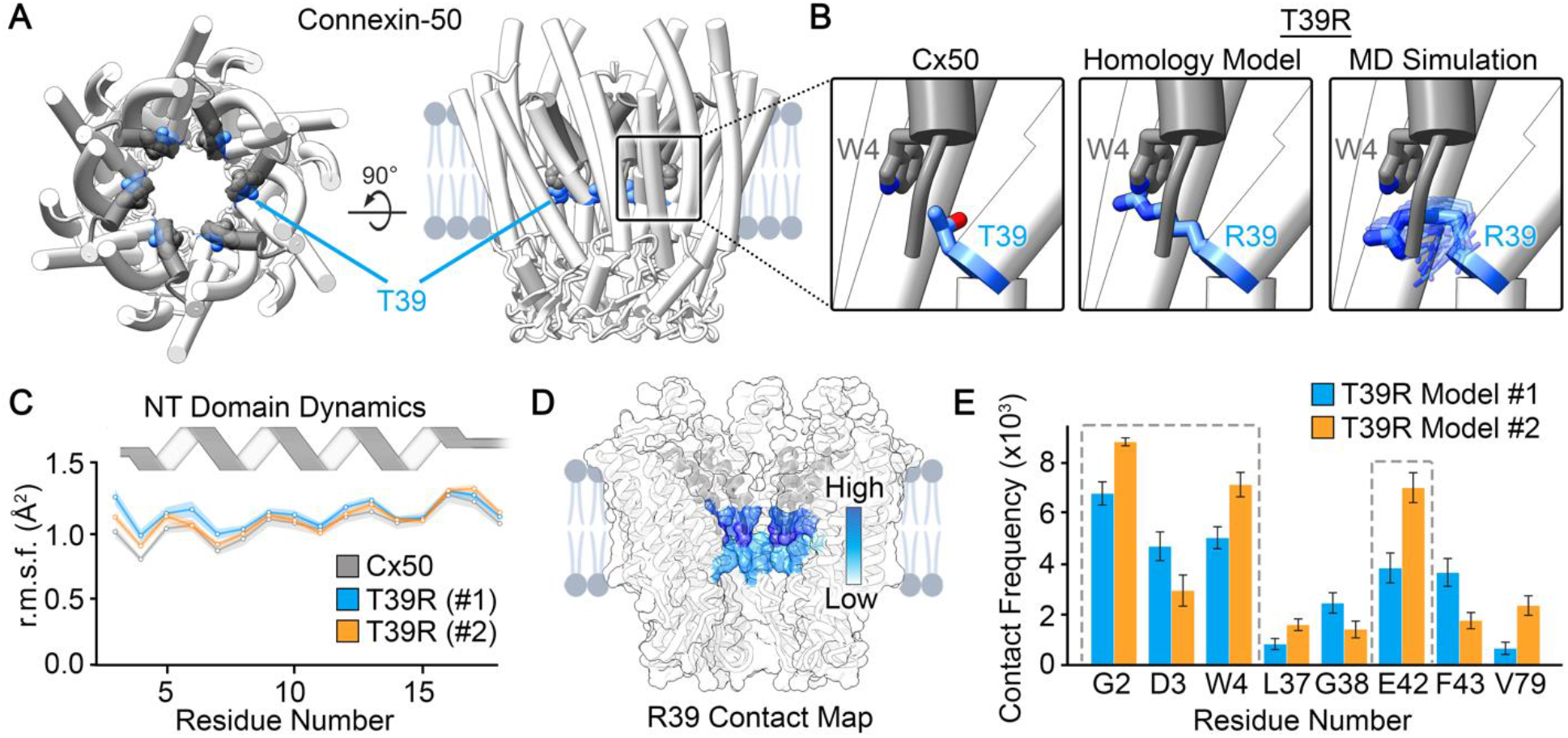
Molecular dynamics simulation of T39R. A. Illustration representing an en face (left) and a side view (right) of a Cx50 hemichannel with the NT domain in grey and T39 in blue (PDB: 7JJP; Flores et al., 2020). B. Magnified views showing that T39 is positioned beneath the NT domain near the hydrophobic anchoring residue W4 in Cx50 (left), the position of T39R based on a naïve homology model (center), and representative poses of R39 following all-atom equilibrium MD simulation (Model #2) (right). C. Root-mean-square-fluctuation (r.m.s.f.) analysis of the NT domain obtained by MD simulation on T39R models (Model #1 – blue; Model #2 – orange) and Cx50 (grey). Solid line indicates the mean with ± SEM in light shading. Each data point represents the average of 12 subunits from each replicate (n = 36 for T39R Model #1 and #2, and n = 24 for Cx50). Grey ribbon diagram illustrates the helical secondary structure of the NT domain. D. Surface representation of Cx50 with the frequency of R39 contacts mapped in color (white = no contact; dark blue = high contact frequency). E. Histogram showing R39 contact frequency by residue number (Model #1 – blue; Model #2 – orange). Error bars represent ± SEM. Grey dashed line boxes indicate regions of high contact frequency that were further investigated.

Contact analyses of R39 highlighted a cluster of amino acids that are within interacting distance to the introduced positively charged guanidinium group. The sites with the highest contact frequency were consistent in both MD simulation models, and localize to the proximal region of the NT domain, and surrounding residues on TM1 (Figure 4D,E). The interaction with the NT domain were localized to residues G2, D3 and W4, while the most significant interaction within TM1 localized to the negatively charged residue E42. Given the role of the NT domain in voltage-sensing and potential coupling between TM1 and the voltage-gating response [29, 40–42, 44–46], we conducted a detailed analysis of these interacting residues.

### T39R forms a salt-bridge interaction with the voltage-sensing residue D3

The residue D3, located at the proximal end of the NT domain, has been shown to function as a voltage-sensor in Cx50 [40, 42] as well as in other connexins [40, 41, 47]. Remarkably, in each of the simulations conducted, the large, positively charged side chain of R39 extended towards the negatively charged carboxylate group on D3, located on a neighboring subunit, forming an apparent salt-bridge interaction (Figure 5A and Supplemental Movie 1). This charge-neutralizing interaction might alter the voltage sensitivity of Cx50T39R. To assess the prevalence of this interaction we conducted a detailed contact distance analysis, which showed that this neutralizing salt-bridge interaction accounts for ∼9 – 21% of the conformational states sampled by R39 during the MD production period and for a significant energetic contribution compared with Cx50, which would increase the energy barrier to move toward the closed state (Figure 5B). Dwell-time analysis of the R39-D3 interaction indicates transient stability, with the majority of interactions lasting < 50 ns, although in a few instances the interaction persisted through the duration of the MD production runs (100 ns) (Figure 5C,D).

**Figure 5.**
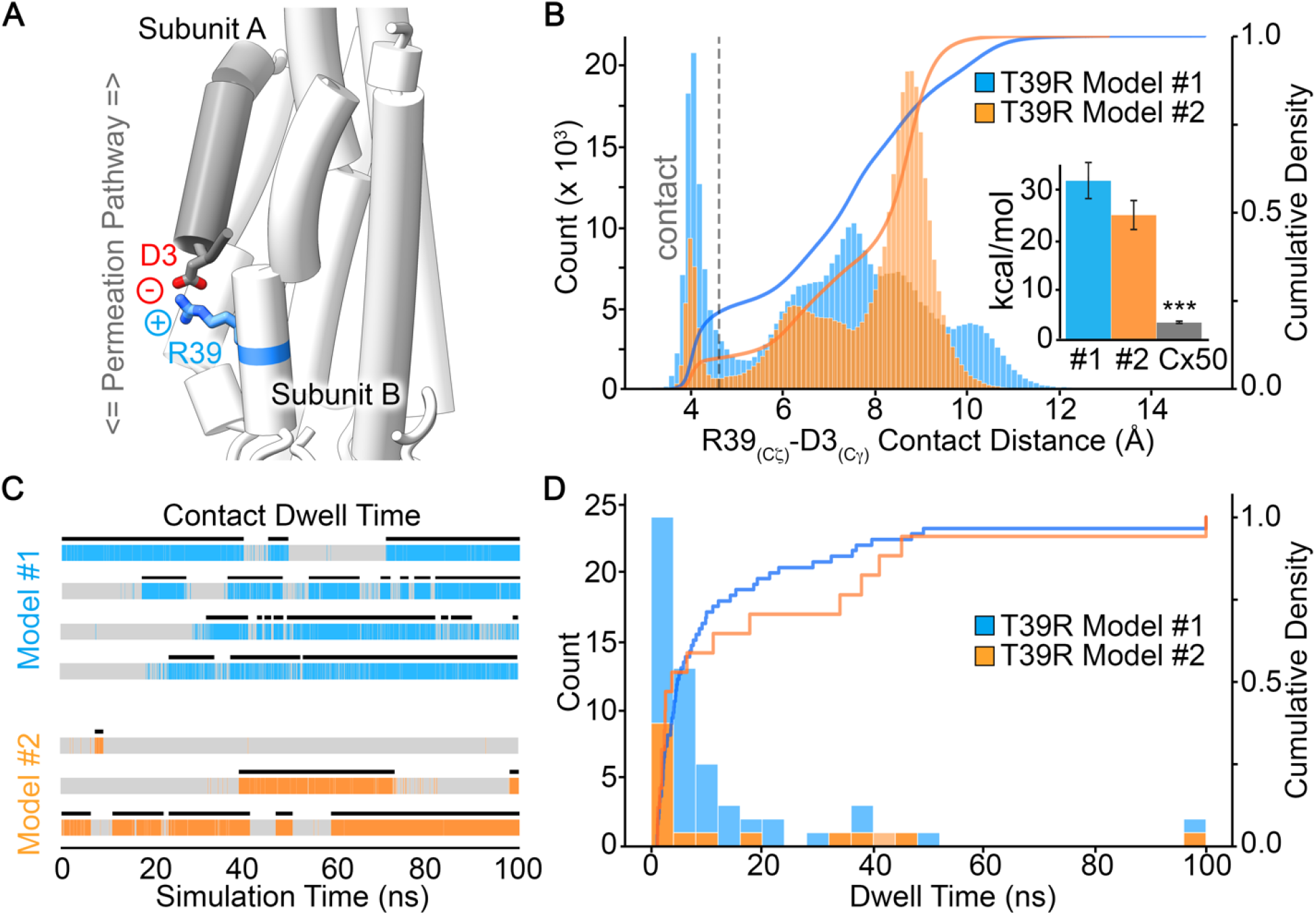
T39R forms a neutralizing salt-bridge interaction with D3. A. Representative snapshot of an MD trajectory illustrating a salt-bridge interaction between the positively charged R39 and the negatively charged residue D3, from a neighboring subunit. B. Histogram showing the population distribution of distance measurements between R39 Cζ and D3 Cγ obtained from the MD trajectories. Distances less than 4.5 Å separating these two heavy atoms are considered to be within contact distance (indicated by the grey dashed line). The cumulative density profile (right axis) is overlaid as a solid line. *Inset,* shows non-bonded energies (kcal/mol) between residue 39 and D3 in the T39R models and Cx50, calculated as the mean ± SEM from each subunit over 3 replicates (n = 36 for T39R Model #1 and #2, and n = 24 for Cx50; *** indicates p value < 0.001). C. R39-D3 dwell-time analysis for representative subunit trajectories, shown in color. Black bars indicate the filtered dwell time in each of the trajectories. D. Histogram showing the population distribution of filtered dwell-time measurements. The cumulative density profile (right axis) is overlaid as a solid line. In panels B-D, data are presented for Model #1 (blue) and Model #2 (orange).

### R39 forms additional interactions with G2 and E42

Given the transient nature of the R39-D3 interaction, we next sought for additional interactions between R39 and the NT domain that might contribute to the drastic increase in open-state stability of T39R observed by electrophysiology. Remarkably, R39 adopts an alternative downward conformation that facilitates simultaneous interaction with the two partial negative charges presented by the carbonyl oxygens on the acetylated G2 residue (G2_ACE_) and also with the negatively charged residue E42 from a neighboring subunit (Figure 6A and Supplemental Movie 2). Such interactions, in particular with G2, could further stabilize the open-state conformation of the NT domain. Contact distance analysis indicates that interactions between R39 and G2_ACE_ are highly populated, accounting for 46 – 62% of the conformational states adopted by R39 during the MD simulation runs (Figure 6B). Similarly, the salt-bridge with E42 was present in 30 – 64% of the conformations sampled in the two MD models (Figure 6C), while no equivalent interaction is made by the wildtype protein (Figure 6C, *inset*). Assessment of non-bonding energies resulting from the R39-G2_ACE_ interaction were significantly greater than in wild-type Cx50 (Figure 6B, *inset*), and consistent with providing additional stability to the open-state conformation of the NT domain. The frequency of these interactions was also significant in both MD models, as assessed by dwell-time analysis (Figure 6D-F). In comparison to the R39-D3 interaction, the coupling to G2 and E42 appears much more frequent and persistent (compare Figure 5D and Figure 6E-F). While the majority of dwell times for the R39-G2_ACE_ and R39-E42 interactions are < 50 ns, a number of trajectories displayed interaction dwell times of 50 – 100 ns (Figure 6E,F).

**Figure 6.**
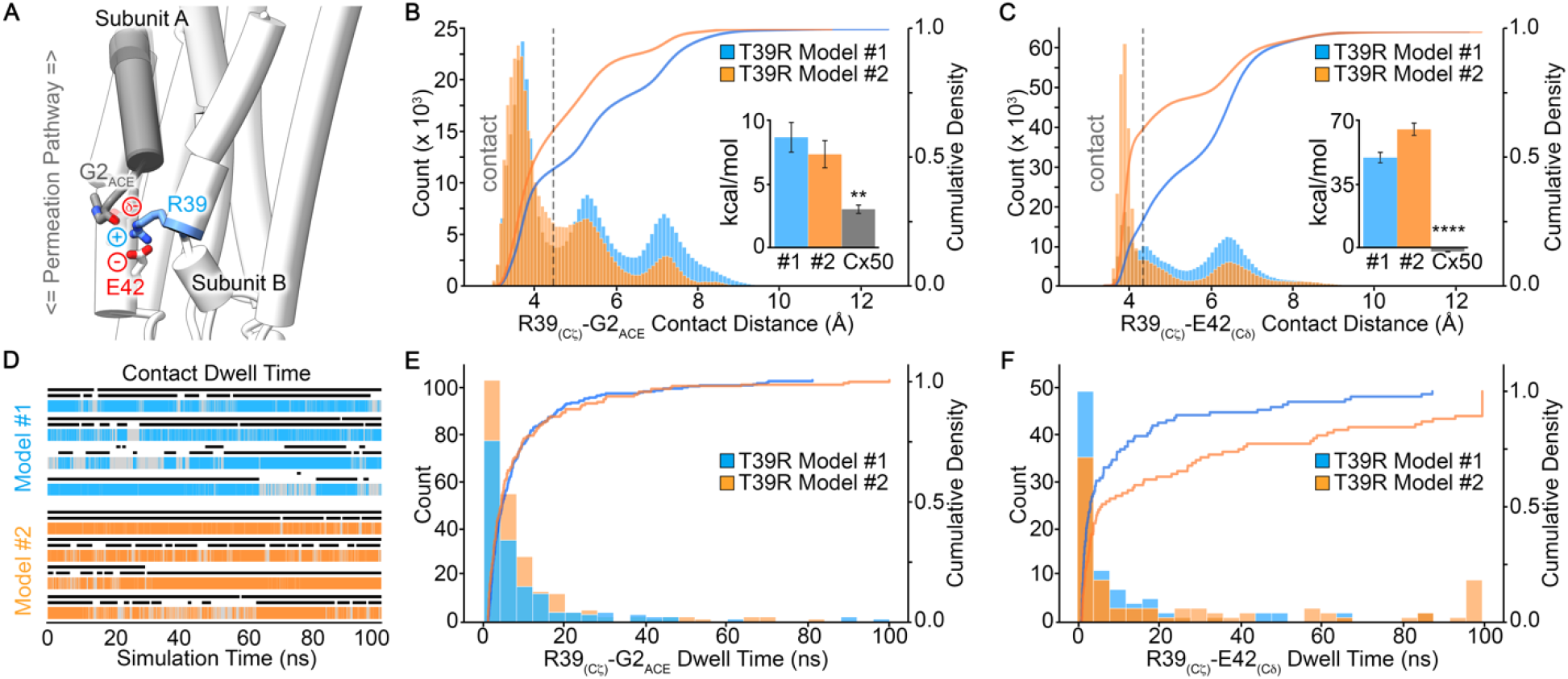
T39R forms a stable interaction with G2_ACE_ and E42. A. Representative snapshot of an MD trajectory illustrating a possible triad interaction between the positively charged R39 and negatively charged residue E42 and acetylated G2 (G2_ACE_), from a neighboring subunit. B and C. Histograms showing the population distribution of distance measurements obtained from the MD trajectories between R39 Cζ and carbonyl oxygens on G2_ACE_ (panel B) and E42 Cδ (panel C). Distances less than 4.5 Å separating heavy atoms are considered to be within contact distance (indicated by the grey, dashed line). The cumulative density profile (right axis) is overlaid as a solid line. *Inset,* in panels B and C shows non-bonded energies (kcal/mol) involving the respective interactions with residue 39 in the T39R models and Cx50, calculated as the mean ± SEM from each subunit over 3 replicates (n = 36 for T39R Model #1 and #2, and n = 24 for Cx50; ** indicates p value < 0.01; **** indicates p value < 0.0001). D. R39-G2_ACE_ dwell-time analysis for representative subunit trajectories, shown in color. Black bars indicate the filtered dwell time of interaction between R39 and E42 (top) and R39 and G2_ACE_ (bottom). E and F. Histogram showing the population distribution of filtered dwell-time measurements for R39-G2_ACE_ (panel E) and R39-E42 (panel F). The cumulative density (right axis) profile is overlaid as a solid line. In panels B-F, data are presented for Model #1 (blue) and Model #2 (orange).

Notably, the conformational state supporting the interaction with G2_ACE_ appears to occur concomitantly with the salt-bridge with E42 (Figure 6A,D). Corresponding dwell-time analysis of the E42 interaction is overlaid with the G2 interaction in Figure 6D (top black bars vs. bottom black bars, respectively). This analysis indicates that R39 interaction with G2 may be supported by the salt-bridge interaction with E42; however, it is not a requirement as analysis of multiple trajectories show that these interactions are not necessarily correlated.

## Discussion

We studied a human Cx50 mutation associated with congenital cataract, T39R. Oocytes injected with T39R cRNA developed greatly enhanced hemichannel currents compared to wild-type Cx50 leading to cell death. Increasing external calcium partially blocked the hemichannel currents and attenuated cell death. When T39R was co-expressed with wildtype lens connexins, it acted in a dominant manner to increase hemichannel activity. Taken together, these findings suggest that T39R causes cataracts through a gain-of-hemichannel function mechanism. Here, the notion is that a hemichannel should reside primarily in the closed state unless it is docked to another hemichannel forming a gap junctional conduit between cells. If an undocked hemichannel has a high probability of opening, it will cause cell death.

T39 (threonine at residue 39) is conserved among all Cx50 orthologues. In contrast, all Cx46 orthologues contain alanine at this position. High resolution structural studies of heteromeric Cx50/Cx46 sheep lens gap junctional channels by cryo-EM show that residue 39 localizes to the first transmembrane pore lining helix (TM1) and anchors the N-terminus (NT) to the inner wall of the channel via hydrophobic interactions with tryptophan 4 (W4), located on the NT domain of a neighboring protomer (Myers et al., 2018). Replacement of threonine at position 39 in Cx50 by positively charged arginine (T39R) appears to stabilize the open state of the undocked hemichannel. It also causes a partial loss of voltage-dependent closure at both positive and negative transmembrane potentials. All-atom MD simulation studies suggest that R39 forms multiple stabilizing interactions with the NT domain, which has been implicated in voltage gating of hemichannels as well as gap junctional channels and contains the negatively charged residue, D3, which has previously been shown to play a critical role in V_j_ gating of Cx50 hemichannels [40, 42]. R39 forms a relatively stable salt-bridge with D3 that would stabilize the NT against the inner wall of the channel when the channel is in the open state and neutralize the negative charge on D3 leading to a loss of V_j_ gating. In addition, R39 adopts an alternative conformational state that facilitates electrostatic interaction with the acetylated G2 position, providing further stability to the open-state conformation of the NT domain. Together, these interactions with the NT domain account for as much as ∼67 – 71% of the conformational states observed for R39 by MD.

Remarkably, we observed that R39 forms an additional neutralizing salt-bridge with E42, as part of a possible tripartite interaction with G2 (see Figure 5). Structurally, E42 is located at the distal end of TM1, near the first extracellular loop, (aka parahelix) and directly adjacent to a Ca^2+^ binding site identified in a crystal structure of Cx26 [48]. This residue contributes to a highly conserved electrostatic network within the extracellular vestibule implicated in hemichannel Ca^2+^ sensitivity and voltage-dependent loop gating in Cx26 and Cx46 [49, 50]. The neutralizing interaction with R39 would be expected to disrupt this electrostatic network and/or Ca^2+^ binding, and therefore potentially further contribute to the attenuated loop gating behavior displayed by T39R.

The role of the positive charge at residue 39 in stabilizing the channel open-state, disabling the voltage-sensing residue D3, and interacting with the negatively charged residue E42 is further supported by experiments replacing threonine at position 39 with alanine (uncharged) or serine (polar). These variants also produced an increase in hemichannel activity albeit to a lesser extent than the arginine variant; however, unlike T39R they retained V_j_ and loop gating. These results are consistent with the inability of these residues to form similar charge-neutralizing salt-bridge interactions with D3 and E42. However, these results also suggest that even small changes at the T39 position can lead to significant alterations in the hydrophobic anchoring interactions of the NT domain with the pore forming TM1/TM2 domain and affect the stability of the open state. We have observed a similar increase in hemichannel activity to occur in a chicken Cx50 chimera in which the first transmembrane domain of Cx50 was replaced by the corresponding domain of Cx46, resulting in the substitution of threonine for alanine at residue 39 [29]. At the same time, we have observed that NT domain swapped chimeras in sheep Cx46 and Cx50 gap junction channels stabilize or destabilize the open-state, depending on the context [46]. Another explanation for the effect of these amino acid substitutions on hemichannel activity is that they may destabilize the closed state. Unfortunately, there are currently no atomic resolution structural models of lens undocked Cx50 hemichannels (or Cx50 gap junctional channels) residing in the closed state. Thus, it is not possible to predict what effect amino acid substitutions at this position might have on the structure of the closed channel.

Our data are also consistent with a dominant gain-of-hemichannel function for T39R. Interactions between wild-type and mutant Cx50 failed to alter the activity or voltage gating properties of T39R hemichannels, suggesting that T39R interacted with Cx50 in a dominant manner. The latter is supported by our MD simulation results, where R39 forms stabilizing interactions with the NT domain of a neighboring subunit. Furthermore, co-expression of T39R with Cx46 resulted in the formation of heteromeric hemichannels that exhibited increased activity at negative potentials, suggesting that they may contribute to the toxic effect of T39R in the lens. In contrast, co-expression of Cx50 with Cx46 was associated with a marked reduction in hemichannel activity indicating that it may have a protective effect.

Comparison of T39R with another Cx50 cataract associated mutant, G46V, showed both similarities and differences [17, 21]. Like T39R, *Xenopus* oocytes expressing G46V developed large, calcium-sensitive hemichannel currents. However, G46V induced smaller transmembrane currents than T39R when expressed in *Xenopus* oocytes at comparable levels and it did not disrupt voltage-dependent closure of Cx50 hemichannels at positive or negative potentials.

In conclusion, our results suggest that T39R causes cataracts by greatly increasing the probability of hemichannel opening in the undocked state. This increase in hemichannel activity would be expected to lead to cell depolarization, loss of ionic gradients, increase in intracellular calcium, and depletion of intracellular metabolites resulting in widespread cell death and the development of total cataract.

## Acknowledgments

This work was supported by the National Institutes of Health (R01-EY026902 to L.E.), (R01-EY030914 to E.C.B. and V.M.B.), (R35-GM124779 to S.L.R.) and (F31-EY031580 to B.G.H.) and the National Science Foundation (XSEDE EMPOWER, ACI-1548562 to U.K.). We are also grateful for access and support at the OHSU Advanced Computing Center.

## Author contributions

All authors contributed to the preparation of the manuscript. J.T. constructed the Cx50 variants, performed the electrophysiology experiments and analyzed the functional data. L.E supervised the electrophysiology studies and analyzed the functional data. U.K. and B.G.H. developed homology structure models, conducted MD simulations and performed analysis of MD data. S.L.R. supervised the MD studies. P.J.M. performed the immunoblot experiments. P.J.M. and V.M.B. analyzed these results. S.L.R. and L.E. provided overall oversight to the design and execution of the work.

## Supplemental Movie Legends

**Supplemental Movie 1. T39R forms a neutralizing salt-bridge interaction with D3.** Representative MD trajectory (100 ns) of T39R forming a salt-bridge between the positively charged R39 and the negatively charged residue D3, from a neighboring subunit

**Supplemental Movie 2. T39R forms a stable interaction with G2_ACE_ and E42.** Representative MD trajectory (100 ns) of T39R forming a triad interaction between the positively charged R39 and negatively charged residue E42 and acetylated G2 (G2_ACE_), from a neighboring subunit.

